# A comparison of survey method efficiency for estimating densities of Zebra Mussels (Dreissena polymorpha)

**DOI:** 10.1101/2022.12.12.520173

**Authors:** Jake M. Ferguson, Laura Jiménez, Aislyn A. Keyes, Austen Hilding, Michael A. McCartney, Katie St. Clair, Douglas H. Johnson, John Fieberg

## Abstract

Abundance surveys are a common and practical technique to estimate plant or animal densities. Methods used to conduct these surveys often require jointly estimating uncertainty in both counts and detection probability. The estimation of detection probability requires additional measurements that take time, potentially reducing the efficiency of the survey for high-density populations. We conducted quadrat, removal, and distance surveys of zebra mussels (Dreissena polymorpha) in three central Minnesota lakes and used an efficiency measure to help compare survey designs. Survey efficiency depended on both the survey design and the population density. The efficiency of survey designs that required estimates of detection probabilities (relative to designs that did not) decreased with density, leading to a change in the most efficient survey design at high densities. These results demonstrate that the best survey design may be context-specific, requiring some prior knowledge of the underlying population density and the cost/time needed to collect additional information for estimating detection probabilities.

## 1 INTRODUCTION

In Ferguson et al. (2019), we explored the application of distance sampling for estimating the abundance of zebra mussels (Dreissena polymorpha) in newly infested lakes. In contrast with conventional distance sampling (Buckland et al., 2001), we found that multiple observers were needed to account for imperfect detection on the transect line. A secondary result was that our observers had significantly different detection probabilities. When discussing these results, one observer reported being more focused on surveying quickly and covering more area while the other observer reported going more slowly to ensure high detection probabilities. This led us to wonder about a potential tradeoff between surveyor speed and detection probability. Specifically, is it better to survey slowly and deliberately, resulting in high detection rates but at the cost of covering less area, or to survey more quickly to cover more area but at the cost of lower detection rates?

Much work has gone into comparing alternative survey designs in specific systems (e.g., Samoilys and Carlos, 2000; Norvell et al., 2003; Pooler and Smith, 2005). Importantly, when choosing between different options, surveyors may be able to implicitly manage tradeoffs between survey coverage and the detectability of targets. At one extreme, basic count methods including quadrat surveys, point counts, and transect surveys are the simplest approaches to implement; however, these methods may require significant effort to ensure every individual is counted, limiting the area that surveyors cover. Furthermore, there are many situations in which it may be impossible to observe all individuals. A variety of methods have been developed to jointly estimate detection and animal density, including mark-recapture, removal (Seber, 1973), distance surveys (Buckland et al., 2001), N-mixture models (Royle, 2004), and sightability surveys (Fieberg, 2012). These approaches may allow a surveyor to move faster and cover more area but at the cost of also needing to account for imperfect detection.

In this study, we examined the efficiency of three survey designs in three lakes that had a range of zebra mussel densities. In each lake, we estimated the density of zebra mussels using quadrat, removal, and distance survey methods. Quadrat surveys are assumed to have perfect detection but may be inefficient at low densities, especially when individuals are clustered (Brown and Manly, 1998), as zebra mussels typically are, due to their association with patchy hard substrate and aggregations of attached individuals known as druses (Karatayev et al., 2015). In distance surveys, observers must measure the perpendicular distance of each detection from the transect line. These distances are then used to model how detection probability declines with distance from the surveyor and to estimate density from the observed counts (Buckland et al., 2001). When densities are low, distance surveys may allow users to cover a larger area in a fixed amount of time relative to quadrat surveys. On the other hand, they may be inefficient at high densities due to the time required to measure the distances between each object and the transect line. Finally, removal surveys (Cook and Jacobson, 1979) utilize a second observer who notes observations missed by a first observer. Removal surveys should require less time than distance surveys to collect the information needed to estimate detection since no distance measurements are required. However, removal surveys will typically have smaller transect widths than distance surveys.

In each lake, we implemented all three survey designs, then compared estimates of zebra mussel density and their associated uncertainties to evaluate the relative performance of the different methods. We predicted that distance surveys would perform best at low densities, removal surveys would perform best at intermediate densities, and quadrat surveys would perform best at high densities.

## 2 METHODS

### 2.1 Field surveys

In Phase 2, we visited six candidate lakes in central Minnesota (Christmas Lake, East Lake Sylvia, Lake Burgan, Lake Florida, Little Birch Lake, and Sylvia Lake). Each of the candidate lakes was recently confirmed by the Minnesota Department of Natural Resources to have zebra mussel infestations. We established 15 survey sites in each lake, approximately evenly distributed around the lake perimeter using ArcMap, then located in the field using a GPS unit (Garmin GPSMAP 64s). Sites occurred at depths between 0.5 to 4.5 m. Based on our past work, we expected that 15 sites would take one to two days to survey for each method, similar to the effort-level that is applied in invasive mussel surveys by local management agencies. At each site, two divers fanned out and spent 20 minutes underwater counting all mussels they encountered. We used these initial timed counts to select three lakes with a range of apparent densities to further survey during the summer of 2018, Lake Florida in Kandiyohi County, Lake Burgan in Douglas County, and Little Birch Lake in Todd County (Figure 1). Lake Florida covers an area of 273 hectares and has a maximum depth of 12 m, Lake Burgan covers an area of 74 hectares and has a maximum depth of 13 m, and Little Birch Lake covers 339 hectares and has a maximum depth of 27 m.

**Figure 1.**
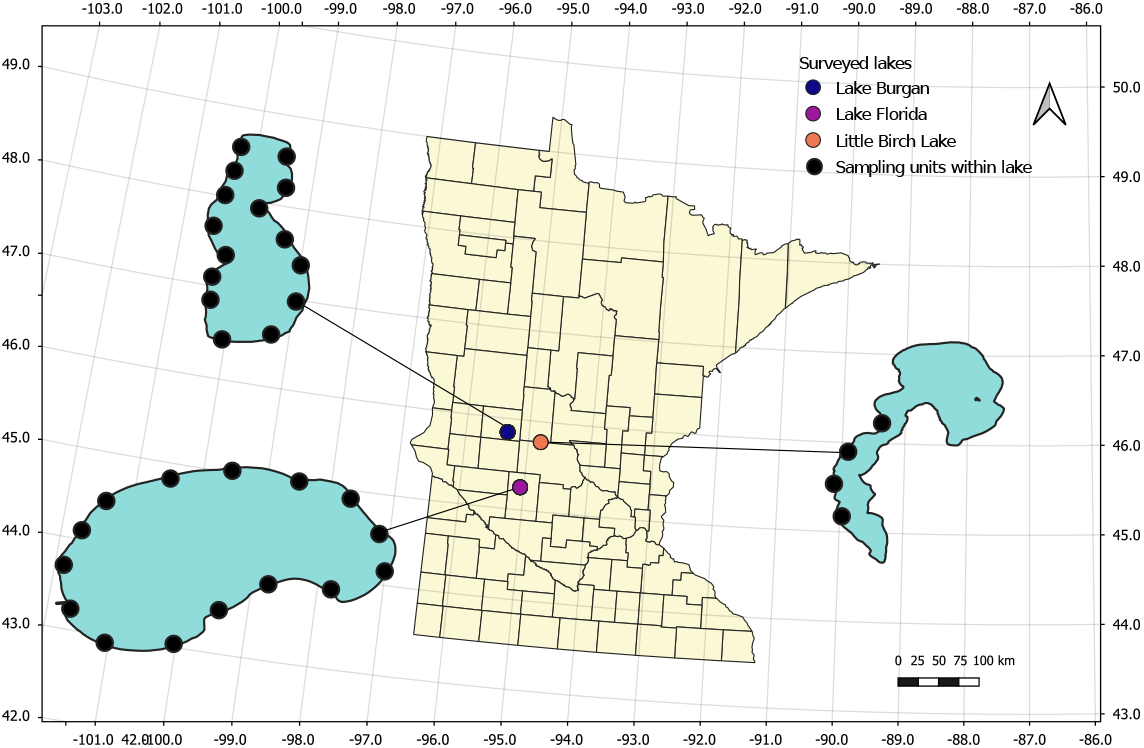
Location of three lakes surveyed in Minnesota during the summer of 2018. Solid black circles within each lake indicate surveyed locations. Lakes are not to scale.

In each of these three lakes, we conducted surveys using three methods: quadrat, removal, and distance-removal surveys. We visited Lake Burgan the second week of July in 2018, followed by Little Birch Lake the next week, and Lake Florida the week after. We spent 3 days surveying each lake, with a single day allocated to each survey method, starting with the removal survey on day one and ending with the quadrat survey on day three. For each survey method, we used the site locations from the Phase 2 timed survey to determine the start of each survey transect. A single day was not sufficient to complete all 15 transects in each lake, so we revisited the lakes in August to complete the remaining transects. We observed high densities of newly-settled zebra mussels in Little Birch Lake from a recruitment event between our first and second visits. Thus, we only used the four transects completed in July when estimating density in Little Birch Lake (Figure 1).

Quadrat surveys were conducted by laying parallel 30-meter transect lines spaced one meter apart at the previously established sites in each lake (Figure 2). Transects were perpendicular to the shoreline. Each diver on our two-person team surveyed one of the parallel transects, placing a 0.5 *×* 0.5 square-meter quadrat every two meters along the transect starting at 0 meters. Divers then counted all zebra mussels within the quadrat. For a full 30-meter transect, this design resulted in 30 quadrat counts (quadrats were not placed at the 30-meter mark since this would result in counting mussels outside of the transect). Both divers sampled an additional 113 quadrats placed along the 11 transects in Little Birch Lake that were not sampled using the other methods due to the recruitment event. Each diver independently sampled these 113 quadrats (hereafter, repeated quadrat counts). The repeated quadrat counts were compared to evaluate the assumption of perfect detection in the quadrat surveys.

**Figure 2.**
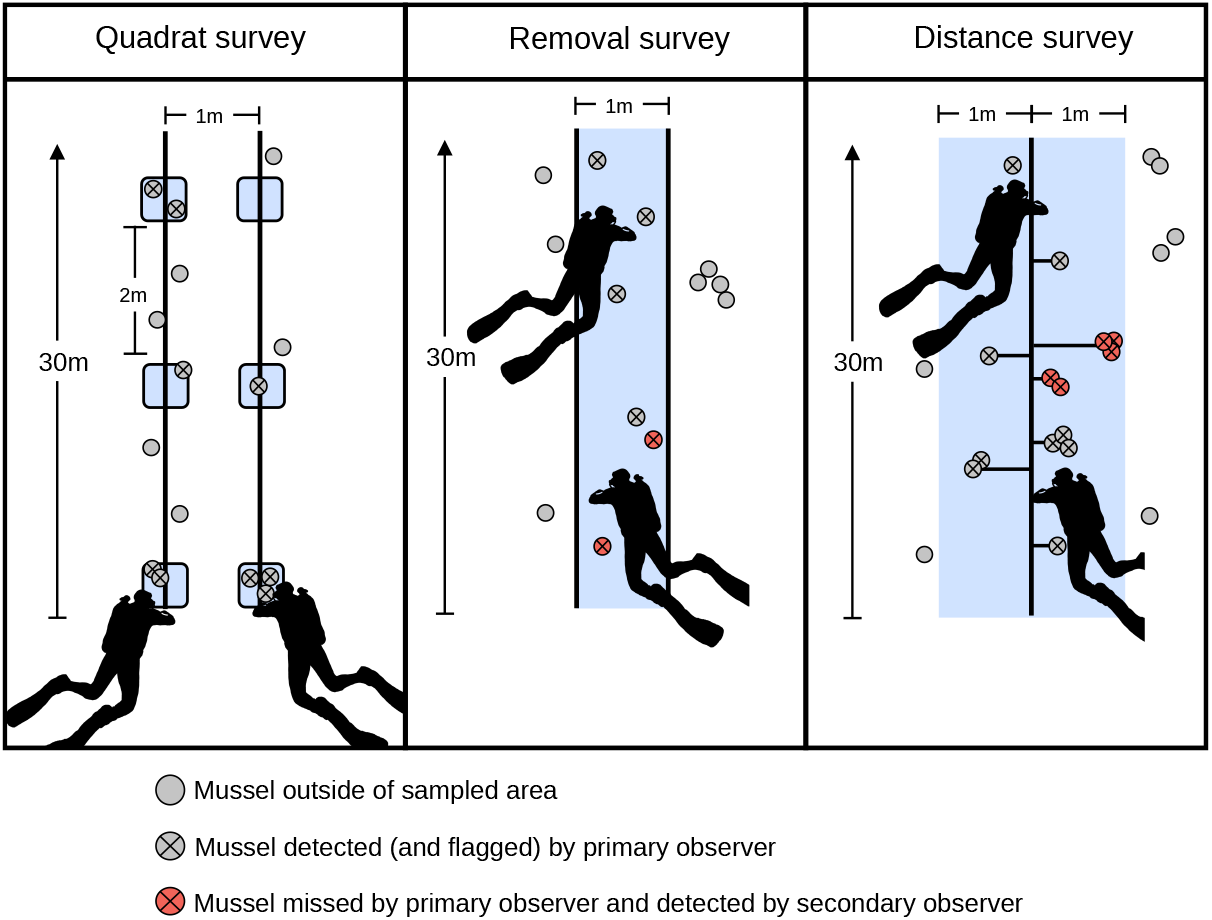
Illustration of a transect for each of the survey techniques used in this study. Blue shaded area indicates the area surveyed by the dive team. Horizontal lines in the distance survey indicate the distance measures used to estimate detection probabilities.

Removal surveys were conducted by laying out parallel 30-meter transect lines spaced one meter apart and perpendicular to the shoreline (Figure 2). These lines allowed divers to determine whether mussels were in the strip-transect. We used the same GPS locations as the quadrat survey to set up the removal transects, although the exact locations of the transects likely differed slightly since we did not physically mark the start of each transect. The first diver swam between the transect lines and whenever the diver detected a zebra mussel or a cluster of mussels, they marked the location with a survey flag and recorded the number of mussels in the cluster, thereby removing the detected individual or cluster from subsequent counts by the second diver. Each cluster was considered a single independent detection event and the number of individual mussels (hereafter cluster size) in the cluster was recorded. The second diver followed behind after a delay to reduce any turbidity that may have been caused by the first diver. They collected the flags and looked for additional mussels missed by the first diver. Divers alternated between the primary and secondary observer roles between transects to average out any innate differences between observers following the recommendation of Cook and Jacobson (1979).

Lastly, we conducted distance-removal (hereafter, distance) surveys. For these, we laid a single 30-meter transect line in a direction perpendicular to the shoreline starting at the same GPS location as the quadrat survey (Figure 2). Divers surveyed up to one meter on either side of the transect. The first diver marked detections, as in the removal survey, then measured the perpendicular distance from the detection to the transect line. The secondary diver then looked for zebra mussels that were missed by the primary diver. We used this approach rather than conventional distance sampling since we previously found that detection along the transect line was imperfect (Ferguson et al., 2019). As in the removal survey, we treated each detected cluster as an independent detection event. For any of the survey techniques, transects were stopped earlier than 30 meters if divers ran into the thermocline due to lowered visibility making counts less reliable.

In addition to the information required to estimate density, we also measured habitat substrate along the transect, proportional plant cover over the length of the transect, and the depth at the start of our transect following methods described in Ferguson et al. (2019), although this information was not used in subsequent analyses. We also recorded the total time required to complete each transect for each survey design and used these measurements to explore differences in the time required to complete surveys by fitting a linear model to the log(transect survey times) with lake, survey method, and their interaction included as predictor variables. The interaction effect allowed us to determine if the relative cost of completing the surveys (measured in units of time) varied across lakes.

#### 2.1.1 Density estimates

Let *n*_*i*_ denote the number of detections for the *i*^th^ transect and 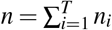 be the total number of detections overall *T* surveyed transects. Further, let *w* denote the transect width, *l*_*i*_ the length of the *i*^th^ transect, and 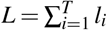 denote the total length of all surveyed transects. We estimated zebra mussel density using (Buckland et al., 2001):

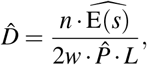

where 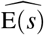 was the estimated average cluster size and 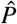 was the estimated average probability of detection. Detection probabilities were obtained using the R package mrds, which takes a model-based approach to estimate detection (Laake et al., 2018). Goodness of fit for the distance models was assessed using a chi-squared test. For both distance and removal surveys, detection probabilities were estimated using maximum likelihood and the variance in the average detection probability was estimated from the Hessian matrix. In the case of quadrat surveys, the detection probability was assumed to be 1 and the variance in the detection probability was assumed to be zero. In the removal survey, we used the removal configuration with full independence, which assumes that observers are independent; in the distance survey, we used the removal configuration coupled with the half-normal detection function for the distance model (code archived at the University of Minnesota Data Repository; temporary link here).

The uncertainty in the estimated density can be approximated by (Buckland et al., 2001):

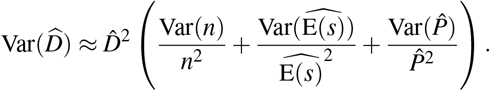

We used a design-based estimator to determine the variance in the counts (Buckland et al., 2001),

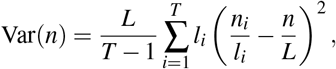

where the contribution of each transect to the total variance was weighted by the transect length, *l*_*i*_. The uncertainty in the average cluster size was calculated as:

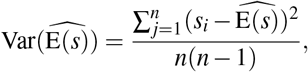

where *s*_*i*_ is the observed cluster-size for the *i*^th^ detection event. For the removal and distance surveys, the uncertainty in 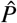 was reported by the mrds package.

We estimated the number of transects needed to achieve a coefficient of variation (hereafter CV) of 0.1 (Buckland, 2006), denoted as *T*_estimated_, by solving the equation 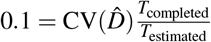, where 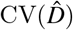 is the CV we obtained from our original survey with *T*_completed_ transects. For Lake Florida and Lake Burgan, *T*_completed_ was 15 transects, whereas *T*_completed_ was 4 for Little Birch Lake (Table 1).

**Table 1.**
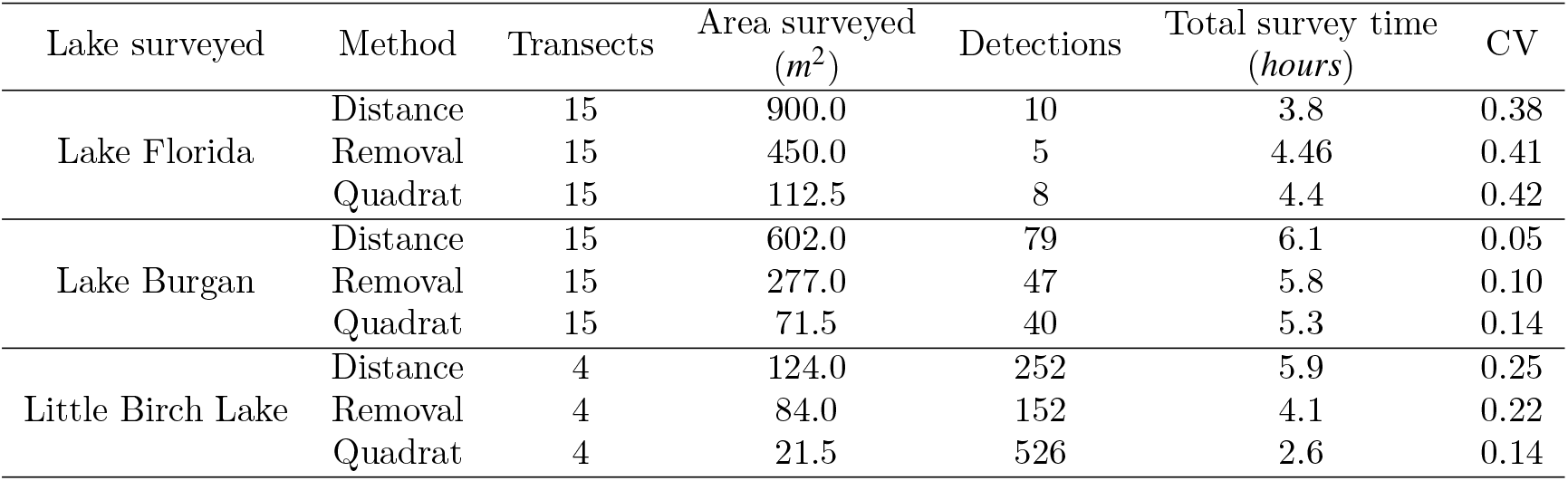
Summary of quadrat, removal, and distance surveys performed in three Central Minnesota lakes surveyed during the summer of 2018. We report the total area surveyed, the number of detected clusters, and the total time taken to complete each survey. CV denotes the coefficient of variation in the estimated density.

## 3 RESULTS

Across the three lakes, we covered the most area using distance surveys (1626 m^2^), compared to 811 m^2^ using removal surveys, and 205.5 m^2^ using quadrat surveys (Table 1). When modeling transect survey times (Figure 3; note the y-axis is log-scaled), the interaction between lake and survey method was statistically significant (*P* = 0.002). In Lake Florida, our lowest density lake, the median transect survey time was similar for removal and quadrat surveys and considerably lower for distance surveys. By contrast, the median transect survey time was lowest for quadrat surveys and highest for distance surveys in our medium- and high-density lakes (Lake Burgan and Little Birch Lake), with removal surveys falling closer to distance surveys than quadrat surveys. Estimated mean transect survey times are provided in Table S2.

**Figure 3.**
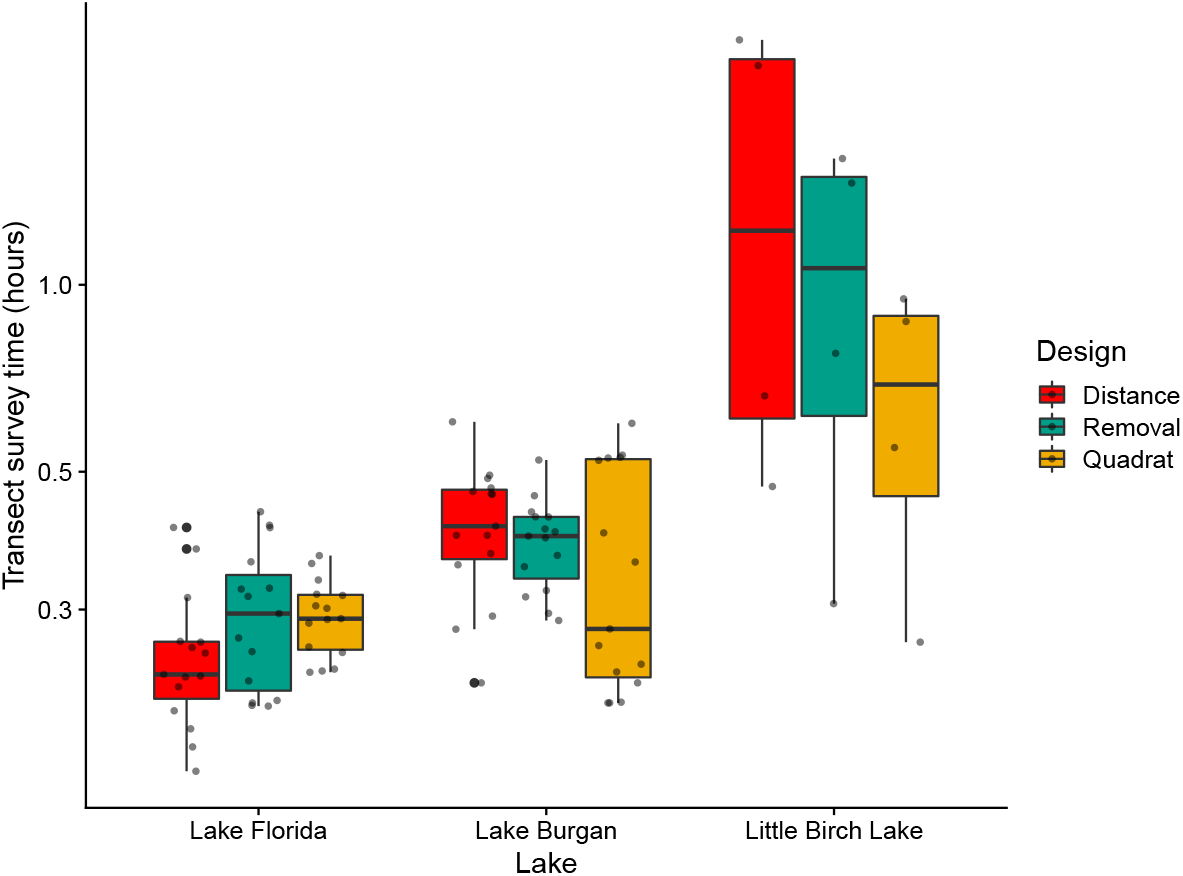
Boxplot indicating the amount of time spent surveying a transect for distance, removal, and quadrat surveys in three Central Minnesota lakes surveyed during the summer of 2018. The lower and upper hinges denote the first and third quartiles, and the horizontal line denotes the median. Points indicate the individual datapoints. The y-axis is log-scaled.

In the repeated quadrat counts in Lake Florida, we found that divers’ counts had observational error (Figure S1); thus, their counts were not perfectly correlated, 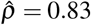. In addition, the amount of error did not appear to vary with the average transect count, suggesting that this error is present even at low densities. While these counts indicated that observational error was present in quadrat counts, subsequent analyses assumed that quadrat counts had no observational error, although we discuss the potential implications of ignoring this observational error in the discussion.

### 3.1 Density estimates

Our estimated probabilities of detection were lower in the distance survey than in the removal survey (Figure 4). The probability of detection in the removal surveys was consistently estimated above 0.9, whereas estimates ranged from about 0.3 to 0.6 in the distance surveys. In the goodness of fit test on the data from Lake Burgan and Little Birch Lake we failed to reject the null hypothesis that our data arose from the half-normal distance function (p-value > 0.05), however, we didn’t have sufficient data to run the test on Lake Florida. In Lake Burgan and Little Birch Lake, the standard errors of the detection probabilities were slightly lower in the removal surveys than the distance surveys, despite removal surveys producing about half the detection events of a distance survey conducted in the same lake (Table 1).

**Figure 4.**
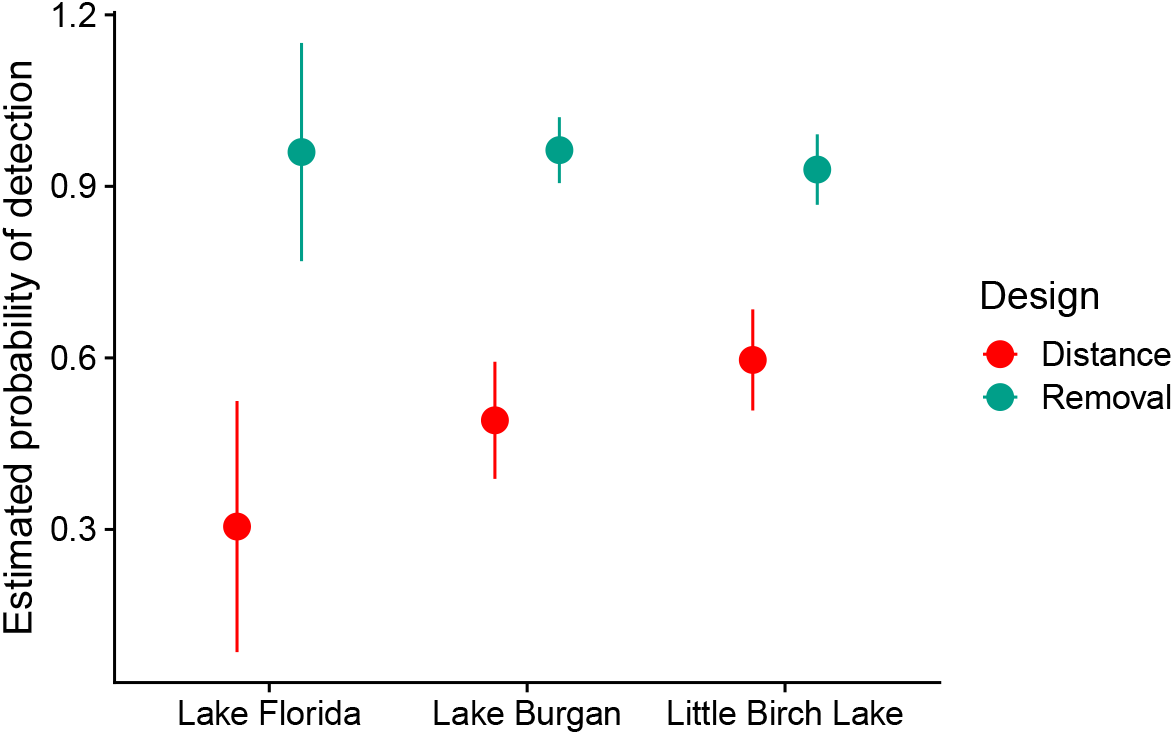
Estimated probability of detection, 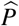, for removal and distance surveys in three Central Minnesota lakes surveyed during the summer of 2018; detection probabilities were assumed to be 1 for quadrat surveys. Error bars denote two standard errors.

Consistent with our initial timed searches on these lakes, we found that Lake Florida had the lowest estimated density, Lake Burgan had an intermediate estimated density, and Little Birch Lake had the highest estimated density (Figure 5). Estimated densities were broadly consistent with other documented invasions in the early phase of their invasion (Strayer et al., 2019). Removal surveys always resulted in the lowest estimated densities. This method assumed that detection probabilities were constant for all mussel clusters. When the true detection probabilities are heterogeneous (e.g., dependent on cluster size or the distance between the cluster and the observer), estimated detection probabilities will be biased high and estimates of density biased low (Caughley and Grice, 1982). This may explain the slightly lower estimates of density using the removal method. Estimates from distance surveys correct for heterogeneity in detection due to distance but could be impacted by heterogeneity associated with cluster size.

**Figure 5.**
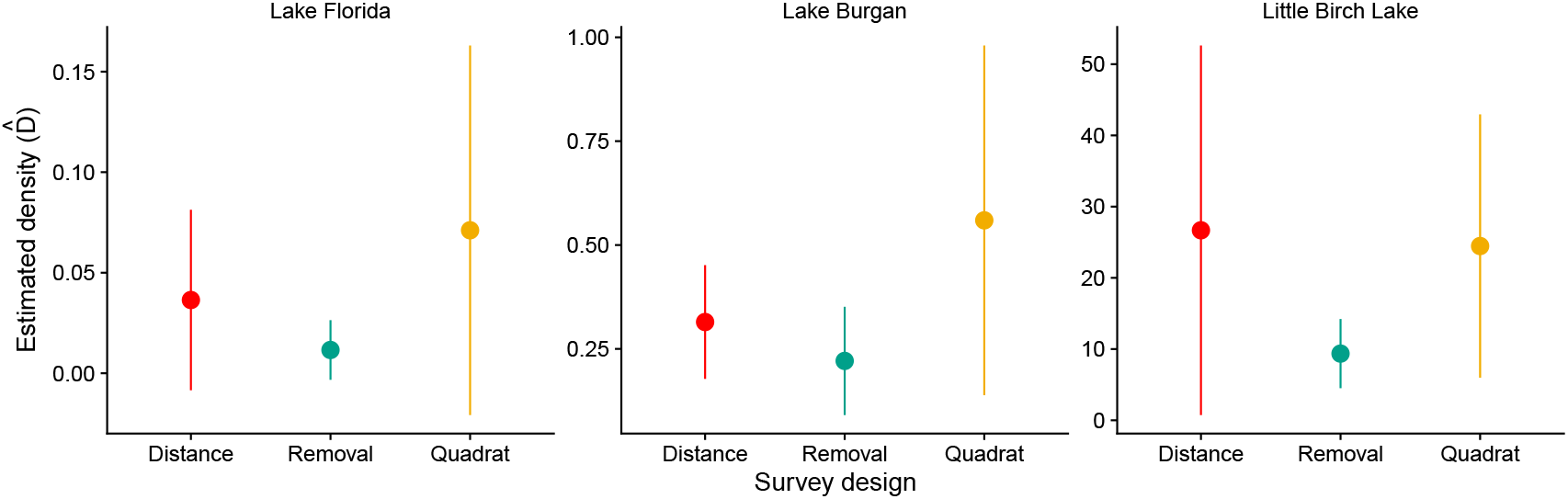
Density estimates (individuals per m^2^) for quadrat, removal, and distance surveys in three Central Minnesota lakes surveyed during the summer of 2018. Error bars denote two standard errors.

The CV in the estimated density was highest for all survey types in Lake Florida for the quadrat survey (Table 1). In the low- and medium-density lakes, the CV was highest in the quadrat surveys, followed by the removal and then the distance surveys. In the highest-density lake, this order was reversed with quadrat surveys having the lowest CV, with removal and distance surveys following.

All mussel detections in Lake Florida were of single zebra mussels, whereas in Lake Burgan, the average cluster size in the removal survey was 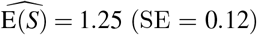 and 1.17 (SE = 0.05) in the distance survey. Clusters were largest in Little Birch Lake, with an average cluster size in the removal survey of 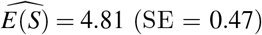 and 7.83 (SE = 0.92) in the distance survey.

The estimated number of transects needed to achieve a CV = 0.1 (Figure 6) was highest in Lake Florida, the low-density lake, require about 60 transects to achieve this goal. The distance survey performed best in this lake (57 transects) followed by the removal survey (61 transects) and quadrat survey (63 transects) (Figure 6). In Lake Burgan and Little Birch Lake, the number of transects needed was less than half that of Lake Florida, indicating that all survey methods were more efficient in higher-density lakes. In Lake Burgan, the distance survey (7 transects) performed better than the removal survey (14 transects) and the quadrat survey (21 transects), and in Little Birch Lake the quadrat survey performed best (6 transects) with the distance (10 transects) and removal (9 transects) surveys performing similarly.

**Figure 6.**
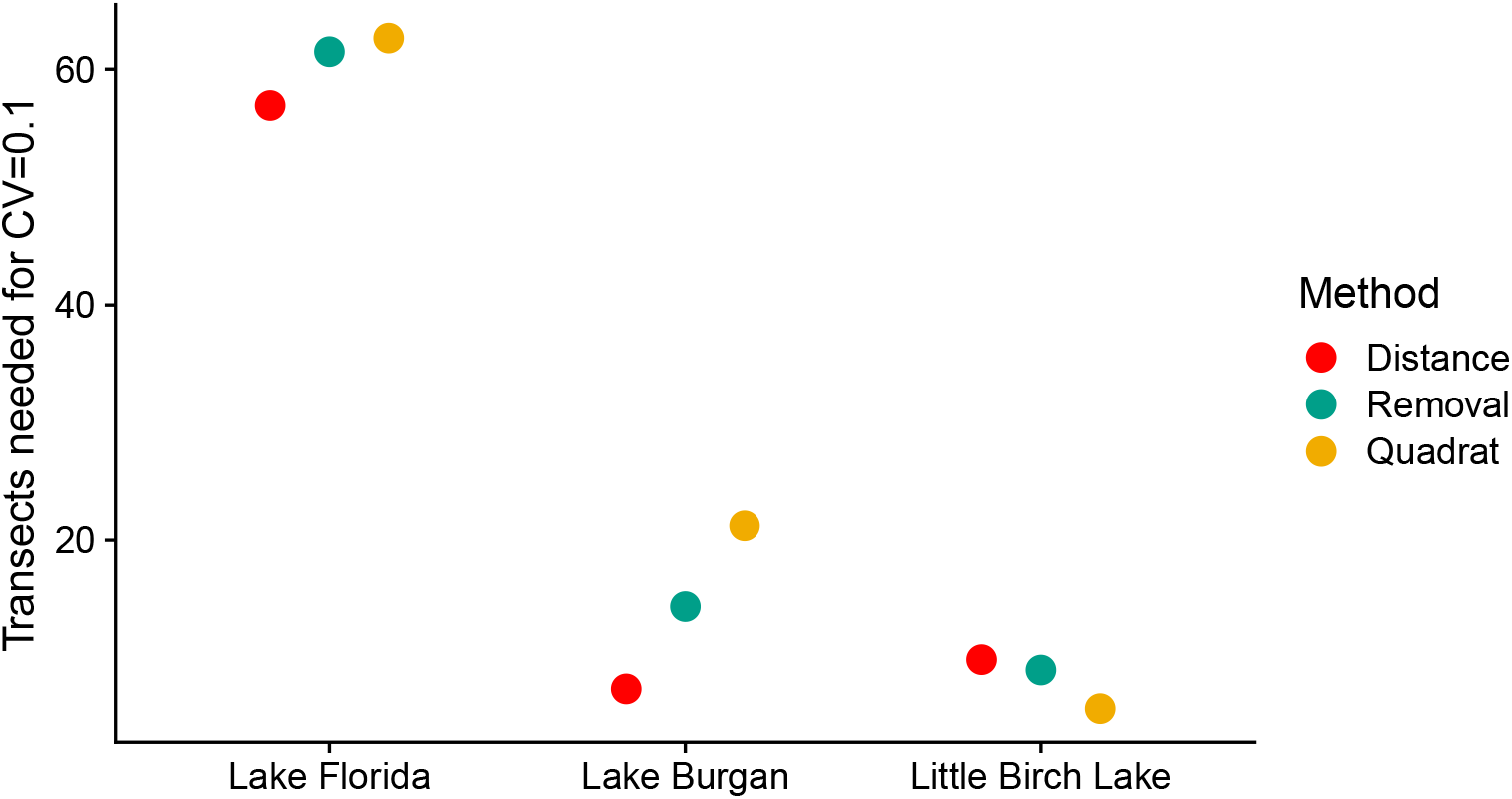
The estimated number of transects needed to achieve a coefficient of variation (CV) of 0.1. Surveys were conducted in three Central Minnesota lakes during the summer of 2018.

Finally, we examined the proportion of the total variance in the estimated density that was due to uncertainty in detection 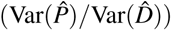. In all lakes, this proportion was small (ranging from 0.01% to 4%), and the contribution was always lower in the removal surveys than in the distance surveys. Thus, uncertainty in detection had a low contribution to the total uncertainty in density.

## 4 DISCUSSION

We found that the relative efficiency of the three survey methods, in terms of the amount of needed effort to achieve a CV = 0.1, varied by lake. In our low- and medium-density lakes, distance removal required the least effort, followed by the removal, then the quadrant surveys. In the high-density lake, this pattern was reversed whereas the quadrat survey was more efficient in our highest-density lake. Distance surveys were more efficient than the removal survey in the low- and medium-density lakes. Broadly, this pattern was consistent with our a priori predictions, though we expected the removal survey to outperform the distance survey in our medium-density lake.

The improved performance of the quadrat survey at the highest densities in our study, relative to the other designs, was likely due to the amount of time it took to record the extra information necessary for estimating detection probabilities. It may be possible to improve the efficiencies of these methods by collecting this information for a subset of transects rather than for all transects (e.g., Pollock et al., 2002). Additional transects could then be completed using a single observer. One caveat of this approach would be that the detection of many animals is known to be habitat-specific (e.g., Aars et al., 2009; Anderson et al., 2015; Ferguson et al., 2019); thus, the transects used to estimate detection should be representative of the available habitat. Work by Knights et al. (2021) has shown that this could be addressed in distance surveys by measuring only a proportion of the targets, where the optimal proportion can be determined when the search time and the time taken to make a detection are known.

One potential advantage of distance surveys is that the transects can be wider than in removal surveys, which could result in more detections and improved estimates of detectability. However, uncertainty in the detection probabilities contributed little to the variance of 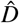, and thus the standard errors were similar for the two methods despite having far fewer detections in the removal survey. Although the distance and removal surveys performed similarly in our study lakes, distance surveys may outperform removal surveys when densities are low and the transect width can be much wider than the width of a comparable removal survey.

Finally, quadrat surveys typically assume perfect detection in each quadrat. Here, we found discrepancies between two observer’s counts of the same quadrat indicating that our counts contained observational error. While we did not account for this error in our estimates, methods do exist to use repeated counts to jointly estimate population size and the probability of detection that could be applied in future surveys (Royle, 2004). The impact of observational error on the density estimates will depend on the details of the detection process. If observers tended to either miss animals or falsely detect animals, we would expect our estimates of density to be biased. However, our quadrat density estimates were consistent with the other survey methods (Figure 5), suggesting that observers were roughly equally likely to over- or under-count the number of individuals. In the presence of both kinds of observational error, we expect the variance in the estimated density to account for both the variance in the counts plus additional variation due to the observational error, leading to higher overall uncertainty in our density estimate.

Our estimated detection probability from the distance survey in Lake Burgan, 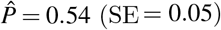 was comparable to the detection probability we estimated for one of our dive teams from the previous field season, 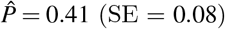 (Ferguson et al., 2019). Yet, it was substantially higher than our estimate of detection for the other dive team, 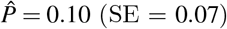 and higher than the average detection probability for the two teams, 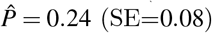 (Ferguson et al., 2019). There are several examples of differences in detection probabilities between observers in surveys of birds (Nichols et al., 2000; Koneff et al., 2008), plants (Moore et al., 2011; Alexander et al., 2012), and mammals (Bauduin et al., 2013; Sunde and Jessen, 2013). This body of work reinforces the importance of accounting for detection when surveys are performed by multiple individuals. Despite the large differences in estimated detection probabilities between years, the estimated densities were remarkably similar, with 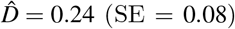 in 2017 and 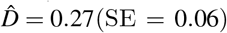 in 2018, indicating that one can obtain consistent estimates when controlling for observer-specific detection probabilities.

Although comparisons among survey methods using empirical data, such as was done in this study, are extremely valuable, they are likely to be highly context-dependent. More general conclusions can sometimes be generated using an analytical framework that captures the costs and efficiencies of different survey methods (e.g., Giudice et al., 2010). We are currently developing a framework that links survey efficiency to measures of effort necessary to perform tasks within each survey (e.g., the amount of time to set up and survey a transect and the additional time required for each detection event). We hope this framework will provide further insights into the tradeoffs between surveyor effort and detection probabilities, and may also prove useful for optimizing this aspect of survey design.

## 5 DATA ACCESSIBILITY

Data and code are available for review here. Data accessibility The data associated with this paper will be made permanently available upon acceptance through Digital Repository of the University of Minnesota at https://doi.org/10.13020/655p-j357.

## 6 ACKNOWLEDGMENTS

This study was funded by the Minnesota Aquatic Invasive Species Research Center with funding from the Minnesota Environmental and Natural Resources Trust Fund as recommended by the Legislative-Citizen Commission on Minnesota Resources. JF received partial support from the Minnesota Agricultural Experimental Station and the McKnight Foundation.

## SUPPLEMENTARY MATERIAL

**Table S1.** Average counts in 30 timed searches (15 minutes each) for six lakes surveyed in Minnesota. Two observers conducted the surveys and visited 15 transects along the perimeter of the lake. Survey results were used to identify three early invaded lakes for subsequent surveys that covered a range of population densities.

| Lake surveyed | Average counts |
| --- | --- |
| Lake Florida | 0.3 |
| Lake Burgan | 1.7 |
| Little Birch Lake | 55.1 |
| East Lake Sylvia | 97.7 |
| Lake Sylvia | 119.5 |
| Christmas Lake | Uncountable |

**Table S2.** Estimated mean transect survey time (in hours) and their associated standard errors.

| Lake | Design | Mean | Standard error |
| --- | --- | --- | --- |
| Lake Burgan | Distance | 0.40 | 0.04 |
|  | Removal | 0.38 | 0.03 |
|  | Quadrat | 0.33 | 0.03 |
| Little Birch Lake | Distance | 1.16 | 0.20 |
|  | Removal | 0.87 | 0.15 |
|  | Quadrat | 0.60 | 0.10 |
| Lake Florida | Distance | 0.25 | 0.02 |
|  | Removal | 0.29 | 0.03 |
|  | Quadrat | 0.29 | 0.03 |

**Figure S1.**
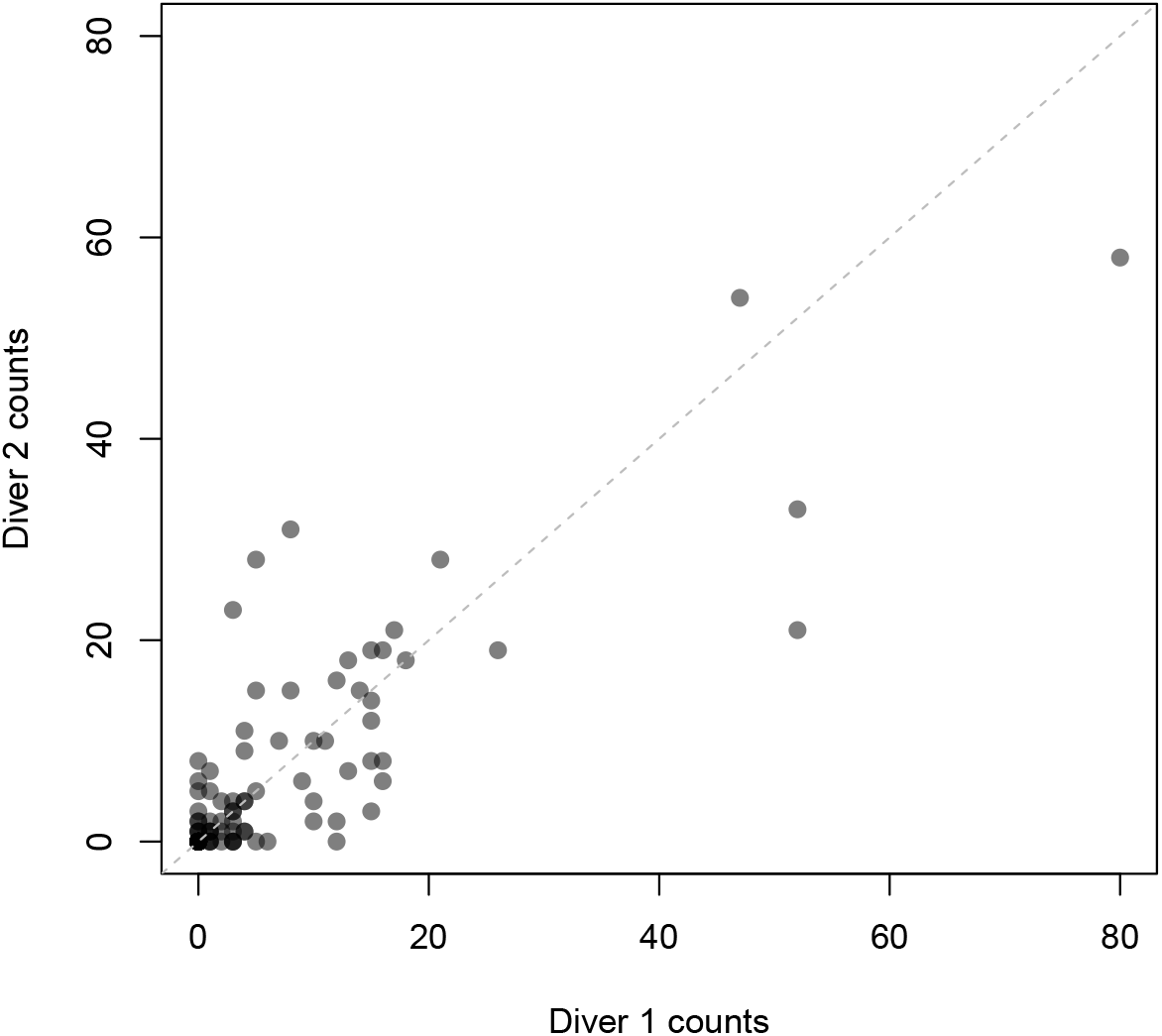
Counts from Diver 1 and Diver 2 on repeated quadrats in 11 transects from Little Birch Lake. Dotted line represents the ideal case where no observational error is present in the counts.

